# Factorial Mendelian randomization: using genetic variants to assess interactions

**DOI:** 10.1101/531228

**Authors:** Jessica MB Rees, Christopher N Foley, Stephen Burgess

## Abstract

**Background:** Factorial Mendelian randomization is the use of genetic variants to answer questions about interactions. Although the approach has been used in applied investigations, little methodological advice is available on how to design or perform a factorial Mendelian randomization analysis. Previous analyses have employed a 2 × 2 approach, using dichotomized genetic scores to divide the population into 4 subgroups as in a factorial randomized trial.

**Methods:** We describe two distinct contexts for factorial Mendelian randomization: investigating interactions between risk factors, and investigating interactions between pharmacological interventions on risk factors. We propose two-stage least squares methods using all available genetic variants and their interactions as instrumental variables, and using continuous genetic scores as instrumental variables rather than dichotomized scores. We illustrate our methods using data from UK Biobank to investigate the interaction between body mass index and alcohol consumption on systolic blood pressure.

**Results:** Simulated and real data show that efficiency is maximized using the full set of interactions between genetic variants as instruments. In the applied example, between four- and ten-fold improvement in efficiency is demonstrated over the 2 × 2 approach. Analyses using continuous genetic scores are more efficient than those using dichotomized scores. Efficiency is improved by finding genetic variants that divide the population at a natural break in the distribution of the risk factor, or else divide the population into more equal sized groups.

**Conclusions:** Previous factorial Mendelian randomization analyses may have been under-powered. Efficiency can be improved by using all genetic variants and their interactions as instrumental variables, rather than the 2 × 2 approach.

**Key messages:**

- Factorial Mendelian randomization is an extension of the Mendelian randomization paradigm to answer questions about interactions.
- There are two contexts in which factorial Mendelian randomization can be used:
for investigating interactions between risk factors, and interactions between
pharmacological interventions on risk factors.
- While most applications of factorial Mendelian randomization have dichotomized the population as in a 2 × 2 factorial randomized trial, this approach is generally inefficient for detecting statistical interactions.
- In the first context, efficiency is maximized by including all genetic variants and their cross-terms as instrumental variables for the two risk factors and their product term.
- In the second context, efficiency is maximized by using continuous genetic scores rather than dichotomized scores.

---

Mendelian randomization is the use of genetic variants as proxies for interventions on risk factors to answer questions of cause and effect using observational data [1, 2]. Formally, Mendelian randomization can be viewed as instrumental variable (IV) analysis using genetic variants as IVs [3, 4]. Factorial Mendelian randomization is the use of genetic variants to answer questions about interactions. It does this by proposing a statistical model for the outcome as a function of the risk factors (or their genetic predictors) and a product term.

A statistical interaction is observed when the coefficient for the product term in the model is non-zero. A statistical interaction in the causal model for the outcome may represent a causal interaction, meaning that the effect of one risk factor on the outcome is dependent upon the value of the other risk factor [5, 6]. This may arise due to a functional or biological interaction, in which there is a mechanistic connection between the two risk factors in how they influence the outcome. However, a statistical interaction may also arise due to non-linearity in the effect of a risk factor, or due to effect modification, in which the effect of one risk factor varies in strata of the other. Hereafter, we take the word ‘interaction’ to mean a statistical interaction in the causal model for the outcome, without implying a functional interaction between the risk factors.

Factorial Mendelian randomization was proposed in the seminal paper on Mendelian randomization by Davey Smith and Ebrahim in 2003 [1]. The term is credited by the authors to Sheila Bird. However, the idea was not readily taken up in applied practice. The concept was raised again by Davey Smith and Hemani in a 2014 review [7], who suggested that genetic predictors of obesity and alcohol consumption could be used to investigate the interaction between the two risk factors on risk of liver disease. This question was investigated for markers of liver function using data from the Copenhagen General Population Study in 2018 [8]; no evidence for an interaction was found.

In parallel to this, the term factorial Mendelian randomization has been used for analyses employing genetic variants as proxies for pharmacological interventions. Ference et al. performed factorial Mendelian randomization to compare the effect of lowering low density lipoprotein (LDL) cholesterol levels on the risk of coronary heart disease (CHD) with two different LDL-cholesterol lowering agents (ezetimibe and statin), and with a combination of both [9]. Genetic variants associated with LDL-cholesterol were identified in the *NPC1L1* gene (proxies for ezetimibe), and the *HMGCR* gene (proxies for statins), and combined into separate gene scores. To mimic a 2 × 2 factorial randomized trial, the two gene scores were dichotomized to create a 2 × 2 contingency table. The gene scores were dichotomized at their median values so that the numbers of participants were balanced across the four groups. Ference has conducted similar analyses for PCSK9 inhibitors and statins [10], and for CETP inhibitors and statins [11]. A similar 2 × 2 approach was used in each case, as well as in the analysis of obesity and alcohol mentioned above [8].

In this paper, we consider various aspects relating to the conceptualization, design, analysis and interpretation of a factorial Mendelian randomization investigation. First, we demonstrate the analogy between factorial Mendelian randomization and a factorial randomized trial, we make a connection with multivariable Mendelian randomization, and we describe two contexts in which factorial Mendelian randomization may have utility: for investigating interactions between risk factors, and for investigating interactions between pharmacological interventions on risk factors. We present simulated data demonstrating that the 2 × 2 approach to analysis, while being conceptually appealing, is inefficient for detecting interactions. The same conclusion is reached in an applied investigation considering interactions between body mass index (BMI) and alcohol consumption on blood pressure using data from UK Biobank. Finally, we discuss the implications of our work to applied factorial Mendelian randomization investigations.

## Methods

### Factorial randomized trials and Mendelian randomization

A factorial randomized trial allows for the simultaneous assessment of two or more treatments in a single study [12]. In its simplest form, a 2 × 2 factorial trial investigates the effect of two binary treatments A and B on a binary outcome *Y*. Participants are randomly allocated to one of four groups: to receive treatment A only; to receive treatment B only; to receive both treatments A and B; or to receive neither treatment A nor B. The analogy between Mendelian randomization and a randomized trial has been made many times [13, 14], and the analogy between factorial Mendelian randomization and a factorial randomized trial has also been made previously in the context of multivariable Mendelian randomization (Figure 1, adapted from [15]).

**Figure 1:**
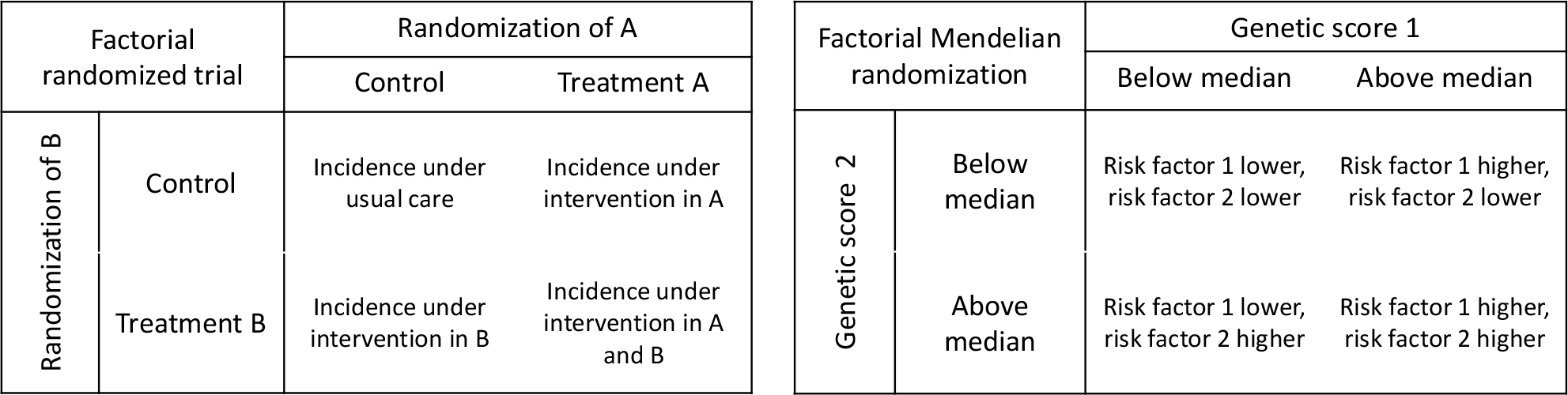
Comparison of a factorial randomized clinical trial and a factorial Mendelian randomization investigation with a 2 × 2 approach (adapted from [15]).

Multivariable Mendelian randomization was motivated by the problem that some genetic variants are associated with multiple risk factors, such that it is impossible to find genetic variants that are specifically associated with a particular risk factor [15]. For illustration, we assume there are two risk factors (*X*_1_ and *X*_2_), and fit a model for the outcome in terms of the risk factors:

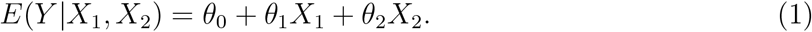

We assume that we have genetic variants *G* that satisfy the multivariable IV assumptions for risk factors *X*_1_ and *X*_2_ [15]. Specifically:

1. Each variant is associated with at least one of the risk factors.
2. Each risk factor is associated with at least one of the genetic variants.
3. Variants are not associated with confounders of the risk factor–outcome
4. Variants are not associated with the outcome conditional on the risk factors and confounders.

If we have at least two genetic variants that are valid multivariable IVs for *X*_1_ and *X*_2_, then causal effects *θ*_1_ and *θ*_2_ can be estimated from the two-stage least squares method by first regressing the risk factors on the genetic variants, and then regressing the outcome on the fitted values of the risk factors from the first-stage regressions [16]. If summarized data on the genetic associations with the outcome 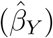 and the risk factors 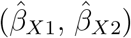 are available, then the same estimates can be obtained by weighted linear regression of the beta-coefficients with the intercept set to zero:

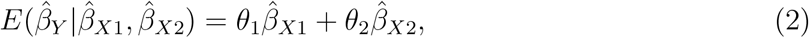

where weights are the reciprocals of variances of the gene–outcome associations 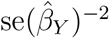[17].

In the language of a factorial randomized trial, this is referred to as an analysis performed ‘at the margins’ [18]. Estimates represent the average direct effect of each of the risk factors [19]. If there is an interaction between the risk factors, then these are marginal estimates – they are averaged over the distribution of the other risk factor.

We can extend multivariable Mendelian randomization by adding a term to the outcome model to estimate an interaction between the risk factors:

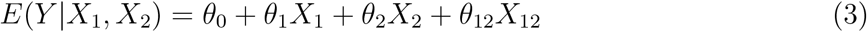

where *X*_12_ is the product *X*_1_ × *X*_2_, and *θ*_12_ is the interaction effect on an additive scale. In a factorial randomized trial, this is referred to as an analysis performed ‘inside the table’, as in a 2 × 2 setting, the interaction can be estimated from the 2 × 2 contingency table [20]. A factorial Mendelian randomization analysis is primarily interested in assessing the presence of, and estimating the interaction effect *θ*_12_.

### Two contexts: interactions between risk factors and interactions between interventions

Factorial Mendelian randomization study has been considered in two broad scenarios: a) to estimate interaction effects between risk factors by using genetic variants as predictors of the risk factors; and b) to identify interactions between interventions by using genetic variants as proxies for specific treatments. In the first case, the aim is to identify an interaction in the effect of two distinct risk factors on the outcome. In the second case, there may not even be two distinct risk factors (as in the example of two LDL-cholesterol lowering interventions discussed by Ference et al. [9]), and the aim is to identify an interaction in the associations of the genetic variants with the outcome. In this case, an interaction is inferred between the interventions for which the genetic variants are proxies. We consider these two scenarios in turn.

### Interactions between risk factors

The multivariable IV assumptions imply that there is no effect of the genetic variants on the outcome except potentially indirectly via one or both of the risk factors. We divide the genetic variants into three groups: *G*_1_ contains variants that are associated with *X*_1_, *G*_2_ contains variants that are associated with *X*_2_, and *G*_*c*_ contains shared variants that are associated with *X*_1_ and *X*_2_ (Figure 2). We can now perform two-stage least squares by first regressing the risk factors *X*_1_, *X*_2_, and the product *X*_12_ on the genetic variants, and then regressing the outcome on the fitted values of these risk factors. This analysis treats *X*_12_ as if it is a separate risk factor unrelated to *X*_1_ and *X*_2_ [21]. For the second-stage regression model to be identified, at least three IVs are required, as three parameters are estimated, and all risk factors (*X*_1_, *X*_2_, *X*_12_) must be associated with at least one IV.

**Figure 2:**
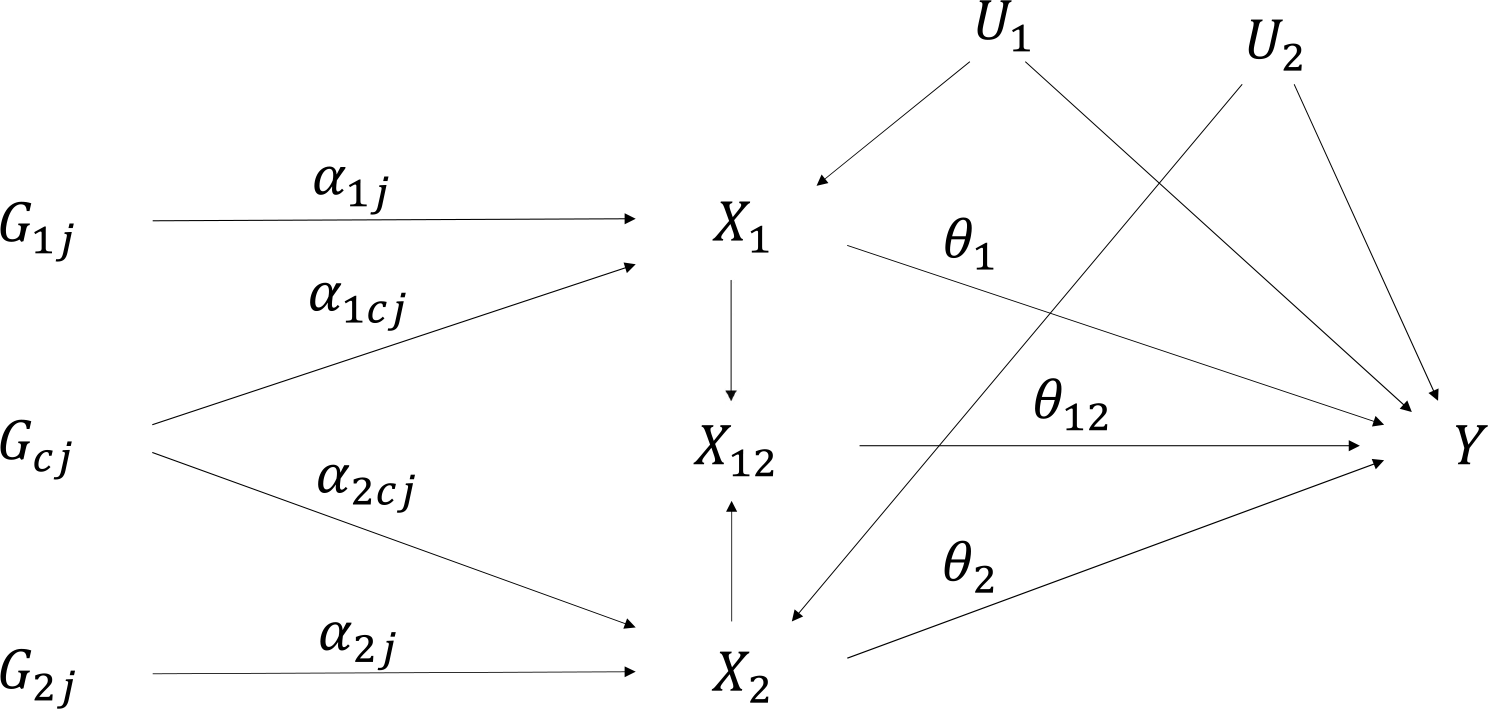
Causal directed acyclic graph illustrating relationships between variables in a factorial Mendelian randomization framework for two risk factors (*X*_1_ and *X*_2_). There are three sets of genetic variants: *G*_1_ (affecting *X*_1_ only), *G*_2_ (affecting *X*_2_ only) and *G*_*c*_ (shared variants, affecting *X*_1_ and *X*_2_). *X*_12_ represents the product *X*_1_ × *X*_2_. The main effects of the risk factors *X*_1_ and *X*_2_ on the outcome *Y* are *θ*_1_ and *θ*_2_, and the interaction effect of *X*_1_ and *X*_2_ on *Y* is *θ*_12_. *U*_1_ and *U*_2_ are sets of confounders.

If we assume that the risk factors *X*_1_ and *X*_2_ are linear in the genetic variants:

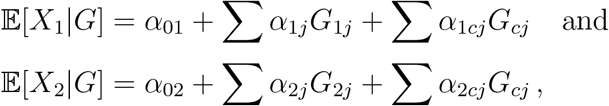

then an interaction between the risk factors means that the statistical model for the outcome includes cross-terms between the genetic variants (such as *G*_11_ × *G*_21_). This motivates the use of cross-terms between the genetic variants as separate IVs.

### Simulation study 1: interactions between risk factors

To investigate the performance of methods for estimating interactions between risk factors, we conduct a simulation study. We assume there are 10 genetic variants that are associated with *X*_1_ and 10 genetic variants that are associated with *X*_2_, and vary the number of shared variants that are associated with both *X*_1_ and *X*_2_ from 0 (20 distinct genetic variants, each associated with one risk factor) to 10 (all 10 genetic variants associated with both risk factors). All genetic variants are simulated as independent (i.e. not in linkage disequilibrium). We compare four methods:

#### Method 1.

Full set of interactions: We consider as IVs all the genetic variants and all cross-terms – so when there are 3 shared variants, there are 114 IVs in total: 7+7+3=17 linear terms, 3 quadratic terms (shared variants only), 3 shared × shared variant cross-terms, 42 shared × non-shared variant cross-terms, and 49 non-shared × non-shared variant cross-terms.

#### Method 2.

Reduced set of interactions: We consider as IVs all the genetic variants and all cross-terms between non-shared variants – so when there are 3 shared variants, there are 17 linear terms and 49 cross-terms.

#### Method 3.

Continuous gene scores: We construct weighted gene scores for each risk factor using external weights, and take the two gene scores and their product as IVs.

#### Method 4.

Dichotomized gene scores: We dichotomize both gene scores at their median, and take the two dichotomized gene scores and their product as IVs. This is equivalent to a 2 × 2 analysis.

The data-generating model for the simulation study is provided in the Supplementary Material. Data were generated 10 000 times for each set of parameters on 10 000 individuals. Parameters were set such that the set of genetic variants explains around 10% of the variance in each risk factor. The effect of *X*_1_ on the outcome was *θ*_1_ = 0.3, the effect of *X*_2_ on the outcome was *θ*_2_ = 0.2, and the interaction effect of *X*_12_ on the outcome took values *θ*_12_ = 0.1, 0.3, and 0.5.

### Simulation study 2: interactions between interventions

We performed a further simulation study to investigate methods for detecting interactions between interventions. We assume there are 3 independent genetic variants that are proxies for intervention A, and the same for intervention B. Fewer variants are considered here as typically variants for such an analysis will come from a single gene region for each risk factor [9]. We compare two approaches:

1. Continuous gene scores: We construct weighted gene scores for changes in the risk factor corresponding to each intervention using external weights, and take the two gene scores and their product as IVs
2. Dichotomized gene scores: We dichotomize both gene scores at their median, and take the two dichotomized gene scores and their product as IVs. This is equivalent to a 2 × 2 analysis.

In each case, we regressed the outcome on the IVs, and estimated an interaction term. As before, the data-generating model for the simulation study is provided in the Supplementary Material. Data were generated 10 000 times for each set of parameters on 10 000 individuals. The interaction effect took values 0.1, 0.3, and 0.5. We varied the minor allele frequencies of the genetic variants used as proxies for interventions A and B, drawing from a uniform distribution between 0.1 and 0.2 (uncommon), or between 0.4 and 0.5 (common), and the proportion of variance explained by the genetic variants (3%, 5% or 7%).

### Applied example: the effects of BMI and alcohol on systolic blood pressure

Increased systolic blood pressure (SBP) is associated with a range of health conditions, including cardiovascular disease and diabetes [22, 23]. Whilst there have been numerous studies highlighting the adverse effects of increased BMI on SBP [24, 25], and the adverse effects of increased alcohol consumption [26], there has been little research on the combined effect of BMI and alcohol consumption on SBP. We illustrate factorial Mendelian randomization by performing an analysis using individual participant data from UK Biobank to estimate the interaction effect of BMI and alcohol consumption on SBP. UK Biobank is a prospective, population-based cohort consisting of around 500,000 participants aged from 40 to 69 years at baseline living in the UK. For the analysis, we considered 291,781 unrelated participants of European descent who passed data quality control measures and had genetic data available.

We used the 77 genome-wide significant variants from a meta-analysis by the Genetic Investigation of ANthropometric Traits (GIANT) consortium in participants of European ancestry to act as IVs for BMI [27]. For alcohol, we identified 10 genetic variants in the *ADH1B* gene region that have been shown to be associated with alcohol consumption [28]. We performed factorial Mendelian randomization analyses using the full set of interactions, continuous gene scores, and dichotomized gene scores. We also performed analyses separately using the lead variant from the *ADH1B* gene region (rs1229984) as the sole IV for alcohol consumption, as was done in the analysis by Carter et al. [8].

## Results

### Simulation study 1: interactions between risk factors

Results from the simulation study for estimating interactions between risk factors are displayed in Table 1 (no shared variants) and Table 2 (varying the number of shared variants). All four approaches provided unbiased estimates of the interaction effect in all scenarios, with coverage for the 95% confidence interval close to the nominal 95% level. Power varied considerably between the methods. With no shared variants, method 1 (full set of interactions) and method 2 (reduced set of interactions) are equivalent and gave the most efficient estimates throughout. Method 3 (continuous gene scores) was less efficient, and method 4 (dichotomized gene scores) was the least efficient. With shared variants, method 1 was the most efficient throughout, and its efficiency was not strongly affected by the risk factors having genetic predictors in common. Between methods 2 and 3, method 2 was more efficient when most of the variants were non-shared, whereas method 3 was more efficient when most of the variants were shared. Again, method 4 was the least efficient in all scenarios. This suggests that the 2 × 2 approach may be underpowered in practice, and instead approaches using all genetic variants and their cross-terms should be considered.

**Table 1.** Simulation study results for interactions between risk factors with no shared variants: median estimate, standard deviation (SD) of estimates, median standard error (SE), empirical power (%) to reject null at 5% significance, and empirical coverage (%) of 95% confidence interval.

|  | Median | SD | Median SE | Power (%) | Coverage (%) |
| --- | --- | --- | --- | --- | --- |
| Methods 1 & 2 <sup>a</sup> – full set of interactions: |  |  |  |  |  |
| $\theta_1 = 0.3$ | 0.3013 | 0.0917 | 0.0910 | 90.2 | 95.0 |
| $\theta_2 = 0.2$ | 0.2022 | 0.0952 | 0.0945 | 57.1 | 94.9 |
| $\theta_{12} = 0.1$ | 0.1101 | 0.0721 | 0.0718 | 33.7 | 94.6 |
| $\theta_1 = 0.3$ | 0.3043 | 0.0918 | 0.0910 | 91.0 | 95.0 |
| $\theta_2 = 0.2$ | 0.2034 | 0.0947 | 0.0945 | 57.9 | 95.5 |
| $\theta_{12} = 0.3$ | 0.3080 | 0.0722 | 0.0718 | 98.8 | 95.2 |
| $\theta_1 = 0.3$ | 0.3048 | 0.0911 | 0.0909 | 90.7 | 95.2 |
| $\theta_2 = 0.2$ | 0.2050 | 0.0944 | 0.0945 | 58.4 | 95.2 |
| $\theta_{12} = 0.5$ | 0.5073 | 0.0715 | 0.0718 | 100.0 | 95.2 |
| Method 3 – continuous gene scores: |  |  |  |  |  |
| $\theta_1 = 0.3$ | 0.2993 | 0.1362 | 0.1333 | 61.4 | 95.4 |
| $\theta_2 = 0.2$ | 0.1991 | 0.1415 | 0.1386 | 30.9 | 95.5 |
| $\theta_{12} = 0.1$ | 0.1010 | 0.1113 | 0.1091 | 15.4 | 95.5 |
| $\theta_1 = 0.3$ | 0.2998 | 0.1359 | 0.1332 | 61.9 | 95.6 |
| $\theta_2 = 0.2$ | 0.2019 | 0.1405 | 0.1387 | 31.5 | 95.8 |
| $\theta_{12} = 0.3$ | 0.3000 | 0.1106 | 0.1091 | 77.5 | 95.8 |
| $\theta_1 = 0.3$ | 0.3004 | 0.1352 | 0.1331 | 61.5 | 95.4 |
| $\theta_2 = 0.2$ | 0.2008 | 0.1409 | 0.1385 | 30.7 | 95.6 |
| $\theta_{12} = 0.5$ | 0.4995 | 0.1107 | 0.1092 | 98.7 | 95.6 |
| Method 4 – dichotomized gene scores: |  |  |  |  |  |
| $\theta_1 = 0.3$ | 0.2986 | 0.2155 | 0.2072 | 31.0 | 95.7 |
| $\theta_2 = 0.2$ | 0.1989 | 0.2246 | 0.2168 | 15.0 | 96.2 |
| $\theta_{12} = 0.1$ | 0.1022 | 0.1786 | 0.1720 | 8.0 | 95.9 |
| $\theta_1 = 0.3$ | 0.3039 | 0.2145 | 0.2074 | 32.1 | 95.8 |
| $\theta_2 = 0.2$ | 0.2047 | 0.2236 | 0.2164 | 15.2 | 96.2 |
| $\theta_{12} = 0.3$ | 0.2972 | 0.1777 | 0.1722 | 41.8 | 96.0 |
| $\theta_1 = 0.3$ | 0.3010 | 0.2148 | 0.2073 | 31.4 | 96.2 |
| $\theta_2 = 0.2$ | 0.2002 | 0.2233 | 0.2163 | 15.3 | 96.1 |
| $\theta_{12} = 0.5$ | 0.5002 | 0.1776 | 0.1718 | 80.7 | 96.1 |

<sup>a</sup>As there are no shared variants, methods 1 and 2 are equivalent.

**Table 2.** Simulation study results for interaction term between risk factors *θ*_12_ = 0.3 varying number of shared variants: median estimate, standard deviation (SD) of estimates, median standard error (SE), empirical power (%) to reject null at 5% significance, and empirical coverage (%) of 95% confidence interval.

| Shared variants | Total IVs | Median | SD | Median SE | Power (%) | Coverage (%) |
| --- | --- | --- | --- | --- | --- | --- |
| Method 1 – full set of interactions: |  |  |  |  |  |  |
| 0 <sup>a</sup> | 120 | 0.3080 | 0.0722 | 0.0718 | 98.8 | 95.2 |
| 1 | 119 | 0.3080 | 0.0723 | 0.0719 | 98.8 | 95.0 |
| 3 | 114 | 0.3090 | 0.0717 | 0.0716 | 98.9 | 95.3 |
| 5 | 105 | 0.3078 | 0.0716 | 0.0707 | 98.9 | 94.9 |
| 8 | 84 | 0.3073 | 0.0682 | 0.0687 | 99.3 | 95.2 |
| 10 | 65 | 0.3056 | 0.0670 | 0.0673 | 99.2 | 95.3 |
| Method 2 – reduced set of interactions: |  |  |  |  |  |  |
| 1 | 100 | 0.3073 | 0.0804 | 0.0794 | 96.7 | 94.9 |
| 3 | 66 | 0.3088 | 0.1003 | 0.0997 | 86.1 | 95.2 |
| 5 | 40 | 0.3056 | 0.1340 | 0.1334 | 63.2 | 95.7 |
| 8 | 16 | 0.3054 | 0.2520 | 0.2471 | 23.9 | 97.1 |
| 10 | 10 | 0.3057 | 0.3883 | 0.3891 | 8.7 | 99.3 |
| Method 3 – continuous gene scores: |  |  |  |  |  |  |
| 0 | 3 | 0.3000 | 0.1106 | 0.1091 | 77.5 | 95.8 |
| 1 | 3 | 0.3005 | 0.1111 | 0.1088 | 77.8 | 95.4 |
| 3 | 3 | 0.2998 | 0.1051 | 0.1048 | 81.0 | 95.6 |
| 5 | 3 | 0.3015 | 0.0997 | 0.0980 | 85.6 | 95.5 |
| 8 | 3 | 0.3003 | 0.0857 | 0.0858 | 93.0 | 95.8 |
| 10 | 3 | 0.2993 | 32.31 | 0.1711 | 42.7 | 99.2 |
| Method 4 – dichotomized gene scores: |  |  |  |  |  |  |
| 0 | 3 | 0.2972 | 0.1777 | 0.1722 | 41.8 | 96.0 |
| 1 | 3 | 0.3028 | 0.1757 | 0.1724 | 42.2 | 96.3 |
| 3 | 3 | 0.3002 | 0.1818 | 0.1773 | 39.8 | 96.4 |
| 5 | 3 | 0.3005 | 0.1948 | 0.1884 | 36.6 | 96.6 |
| 8 | 3 | 0.3007 | 0.2474 | 0.2340 | 25.7 | 97.2 |
| 10 | 3 | 0.2896 | 133.5 | 1.3578 | 0.7 | 100.0 |

<sup>a</sup>When there are no shared variants, methods 1 and 2 are equivalent.

We also varied the strength of the genetic variants due to potential concerns about weak instruments [29]. We considered scenarios in which the genetic variants explained 1% and 5% of variance in the risk factors. Although substantial weak instrument bias was observed for the main effects, no bias was observed for the interaction term, even when there were 100 IVs in the analysis and F-statistics and conditional F-statistics [30] for the product term were around 1 (Supplementary Tables A2 and A3). Similar findings were observed in a one-sample setting when varying the direction of confounder effects on the risk factor and outcome (results not shown). We also performed the simulation study centering the values of the risk factors to reduce the impact of collinearity. This changed the mean estimates of the main effects *θ*_1_ and *θ*_2_ and improved precision for the main effect estimates, but estimates and inferences for the interaction term *θ*_12_ were unchanged (Supplementary Table A1). These additional simulations suggest that factorial Mendelian randomization should only be used when the interaction is the main object of interest, and numerical estimates for the main effects from this model should be interpreted with caution.

### Simulation study 2: interactions between interventions

Results from the simulation study for estimating interactions between interventions are displayed in Table 3. While the numerical values of estimates differed between the two approaches, a consistent finding was that power to detect an interaction was greater using continuous gene scores than using dichotomized gene scores. Varying the proportion of variance explained by the genetic variants had no discernable effect on the power to detect an interaction. This can be seen by comparing scenarios 1, 2, and 3, and scenarios 5 and 6. However, varying the minor allele frequency had a strong effect on power, with greater power when the minor allele frequency was close to 0.5. This can be seen by comparing scenarios 2, 4, and 5, and scenarios 3 and 6. This suggests that ensuring comparable size between subgroups is an important factor for efficient detection of interactions, and can be more important than ensuring that the strongest variant is used in the analysis.

**Table 3.** Simulation study results for interaction between interventions: median estimate, standard deviation (SD) of estimates, median standard error (SE), and empirical power (%) to reject null at 5% significance. The minor allele frequencies and proportion of variance explained for variants that are proxies for interventions A and B are varied between scenarios.

|  | Continuous gene scores |  |  |  | Dichotomized gene scores |  |  |  |
| --- | --- | --- | --- | --- | --- | --- | --- | --- |
|  | Median | SD | Median SE | Power | Median | SD | Median SE | Power |
| Scenario 1: (A) Common variants, 3%; (B) Common variants, 3% |  |  |  |  |  |  |  |  |
| $\theta_{12}=0.1$ | 0.0583 | 0.0420 | 0.0417 | 29.3 | 0.0368 | 0.0423 | 0.0421 | 13.5 |
| $\theta_{12}=0.3$ | 0.0330 | 0.0080 | 0.0078 | 98.7 | 0.1102 | 0.0429 | 0.0423 | 73.5 |
| $\theta_{12}=0.5$ | 0.0224 | 0.0034 | 0.0032 | 100.0 | 0.1846 | 0.0428 | 0.0427 | 98.9 |
| Scenario 2: (A) Common variants, 5%; (B) Common variants, 5% |  |  |  |  |  |  |  |  |
| $\theta_{12}=0.1$ | 0.0484 | 0.0343 | 0.0343 | 29.1 | 0.0372 | 0.0420 | 0.0422 | 13.5 |
| $\theta_{12}=0.3$ | 0.0304 | 0.0074 | 0.0072 | 98.8 | 0.1108 | 0.0424 | 0.0423 | 74.3 |
| $\theta_{12}=0.5$ | 0.0212 | 0.0033 | 0.0030 | 100.0 | 0.1851 | 0.0439 | 0.0427 | 99.0 |
| Scenario 3: (A) Common variants, 3%; (B) Common variants, 7% |  |  |  |  |  |  |  |  |
| $\theta_{12}=0.1$ | 0.0498 | 0.0350 | 0.0352 | 29.2 | 0.0371 | 0.0422 | 0.0422 | 14.1 |
| $\theta_{12}=0.3$ | 0.0305 | 0.0075 | 0.0072 | 99.0 | 0.1106 | 0.0426 | 0.0423 | 74.2 |
| $\theta_{12}=0.5$ | 0.0213 | 0.0033 | 0.0030 | 100.0 | 0.1844 | 0.0430 | 0.0427 | 99.1 |
| Scenario 4: (A) Uncommon variants, 5%; (B) Uncommon variants, 5% |  |  |  |  |  |  |  |  |
| $\theta_{12}=0.1$ | 0.0824 | 0.1152 | 0.1150 | 10.9 | 0.0168 | 0.0435 | 0.0430 | 7.0 |
| $\theta_{12}=0.3$ | 0.1082 | 0.0519 | 0.0500 | 58.8 | 0.0526 | 0.0434 | 0.0430 | 23.3 |
| $\theta_{12}=0.5$ | 0.0996 | 0.0300 | 0.0278 | 94.6 | 0.0879 | 0.0436 | 0.0430 | 53.0 |
| Scenario 5: (A) Common variants, 5%; (B) Uncommon variants, 5% |  |  |  |  |  |  |  |  |
| $\theta_{12}=0.1$ | 0.0669 | 0.0699 | 0.0685 | 16.7 | 0.0246 | 0.0434 | 0.0425 | 9.1 |
| $\theta_{12}=0.3$ | 0.0618 | 0.0211 | 0.0204 | 85.5 | 0.0763 | 0.0433 | 0.0426 | 42.8 |
| $\theta_{12}=0.5$ | 0.0489 | 0.0109 | 0.0097 | 99.9 | 0.1279 | 0.0434 | 0.0428 | 84.1 |
| Scenario 6: (A) Common variants, 3%; (B) Uncommon variants, 7% |  |  |  |  |  |  |  |  |
| $\theta_{12}=0.1$ | 0.0748 | 0.0756 | 0.0742 | 17.8 | 0.0259 | 0.0432 | 0.0426 | 9.7 |
| $\theta_{12}=0.3$ | 0.0649 | 0.0221 | 0.0215 | 85.4 | 0.0758 | 0.0430 | 0.0426 | 42.9 |
| $\theta_{12}=0.5$ | 0.0510 | 0.0113 | 0.0101 | 99.9 | 0.1271 | 0.0435 | 0.0428 | 83.9 |

### Applied example: the effects of BMI and alcohol on systolic blood pressure

The lead variant (rs1229984) explained 0.24% of the variance in alcohol consumption, whereas the 10 variants explained 0.28% of the variance. Although the alcohol-decreasing allele of the rs1229984 variant is dominant, its frequency is only 2.5%. Dichotomizing participants based on this variant led to unequal groups in the population, whereas dichotomizing based on the 10 variant score led to equal groups (Table 4). However, the difference in mean alcohol levels between subgroups was reduced when using the 10 variant score, as most of the difference is due to the rs1229984 variant.

**Table 4.** Numbers (%) of participants and mean (standard deviation) of body mass index, alcohol consumption and systolic blood pressure in 2 × 2 subgroups when either 10 genetic variants or the rs1229984 variant used as IVs for alcohol consumption.

|  | Participants (%) | Mean (SD) |  |  |
| --- | --- | --- | --- | --- |
|  |  | BMI (kg/m<sup>2</sup>) | Alcohol (units/day) | SBP (mmHg) |
| Overall | 291,781 (100.0) | 27.1 (4.51) | 2.54 (2.58) | 140.0 (19.8) |
| 10 variants for alcohol: |  |  |  |  |
| Low BMI, low alcohol | 73,003 (25.0) | 26.6 (4.25) | 2.50 (2.52) | 140.6 (20.6) |
| High BMI, low alcohol | 72,889 (25.0) | 27.5 (4.65) | 2.47 (2.50) | 141.2 (20.6) |
| Low BMI, high alcohol | 72,888 (25.0) | 26.7 (4.30) | 2.61 (2.68) | 140.8 (20.7) |
| High BMI, high alcohol | 73,001 (25.0) | 27.6 (4.71) | 2.59 (2.59) | 141.3 (20.6) |
| rs1229984 variant for alcohol: |  |  |  |  |
| Low BMI, low alcohol | 6,997 (2.4) | 26.3 (4.10) | 2.00 (2.04) | 139.2 (20.2) |
| High BMI, low alcohol | 6,863 (2.4) | 27.3 (4.50) | 1.95 (1.99) | 139.7 (20.2) |
| Low BMI, high alcohol | 138,894 (47.6) | 26.7 (4.28) | 2.59 (2.59) | 140.8 (20.6) |
| High BMI, high alcohol | 139,027 (47.6) | 27.6 (4.69) | 2.56 (2.56) | 141.3 (20.6) |

Estimates of the interaction between BMI and alcohol consumption are displayed in Table 5. For the dichotomized gene scores, efficiency is greater when the rs1229984 variant is used, suggesting the importance of dichotomizing the risk factor at a natural break in its distribution (if one exists) rather than ensuring that subgroups are equal in size. However, efficiency is strikingly improved using the full set of interactions, with the standard error decreasing over ten-fold using the 10 variants, and by a factor of four using the rs1229984 variant, compared to the 2 × 2 analysis. All estimates are compatible with the null, suggesting a lack of interaction in the effects of BMI and alcohol on SBP. There was no evidence of weak instrument bias, even though up to 857 IVs were used in the analyses and F-statistics were generally low (Supplementary Table A4).

**Table 5.** Factorial Mendelian randomization results for applied example: estimate of interaction between BMI and alcohol consumption on systolic blood pressure. Estimates are in mmHg units per 1 kg/m^2^ change in BMI and 1 unit/day change in alcohol consumption.

|  | Total IVs | Estimate | Standard error | p-value |
| --- | --- | --- | --- | --- |
| 10 variants for alcohol |  |  |  |  |
| Method 1: full set of interactions | 857 | 0.0023 | 0.0503 | 0.96 |
| Method 2: continuous gene scores | 3 | 0.0655 | 0.3402 | 0.85 |
| Method 3: binary gene scores | 3 | 0.1011 | 0.6411 | 0.87 |
| rs1229984 variant for alcohol |  |  |  |  |
| Method 1: full set of interactions | 149 | -0.0170 | 0.1136 | 0.88 |
| Method 2: continuous gene scores | 3 | 0.1917 | 0.3725 | 0.61 |
| Method 3: binary gene scores | 3 | 0.1499 | 0.4174 | 0.72 |

## Discussion

In this paper, we have provided a brief review of factorial Mendelian randomization, an approach that uses genetic variants as instrumental variables to detect interactions. We have described two broad scenarios in which factorial Mendelian randomization has been implemented: to explore interactions between risk factors, and to explore interactions between interventions. Although most (perhaps even all) factorial Mendelian randomization analyses have been conducted using a 2×2 approach in which the sample is divided into 4 subgroups, we have shown that this approach is generally inefficient, particularly for exploring interactions between risk factors. This has been demonstrated in simulation studies, and in an applied example in which a four- to ten-fold improvement in efficiency was observed by an analysis using the full set of interactions between the genetic variants as IVs.

### Choice of variants

Our findings suggest that factorial Mendelian randomization analyses should be conducted using all available genetic variants that are valid instruments: that is, that satisfy the multivariable IV assumptions. Analyses should not only include the genetic variants as main effects, but also all relevant two-way cross-terms. If investigators want to perform a 2 × 2 analysis, this should be done to illustrate the method rather than being the main analysis for testing the presence of an interaction. For a 2 × 2 analysis, the primary consideration for choosing genetic variants should be to divide the population at a natural break in the distribution of the risk factor, in order to maximize the difference between the mean level of the risk factor in the two halves of the population. If there is no natural break in the distribution, then investigators should find a division that splits the population as far as possible into equal groups. This may entail selecting genetic variants that explain less variance in the risk factor, but have minor allele frequency closer to 50%. There can also be substantial benefit in including multiple variants in a single gene region in an analysis, even if these variants only explain a small additional proportion of variance in the risk factor.

### Weak instrument bias and efficiency

Conventionally, it is discouraged to use large numbers of genetic variants that are not strongly associated with the risk factor in a Mendelian randomization analysis due to weak instrument bias [31]. Although we did not detect any bias from weak instruments on interaction terms in our simulations, we acknowledge that users of the method may be reluctant to use hundreds of cross-terms as IVs. We would therefore encourage the use of continuous gene score methods as sensitivity analyses. Such analyses estimate fewer parameters, so should be less susceptible to bias. However, this advice is precautionary; no evidence of weak instrument bias in interaction estimates was observed in our simulations.

### Summarized data

While multivariable Mendelian randomization can be performed using summarized data that are typically reported from genome-wide association studies by large consortia, this is not possible for factorial Mendelian randomization. If summarized association estimates are available on genetic associations with the product of the two risk factors, as well as associations with the risk factors individually, then the interaction effect can in principle be estimated by weighted linear regression of the beta-coefficients as in multivariable Mendelian randomization. However, if association estimates are only available for genetic variants, then the regression model is not identified asymptotically due to collinearity, and finite-sample estimates will be biased [32]. Association estimates for some cross-terms of genetic variants are additionally required. Hence, factorial Mendelian randomization can be performed using summarized data, but only if bespoke summarized data are available on associations of genetic variants and their cross-terms with the risk factors and their product.

### Interpretation of the interaction effect

If genetic variants each satisfy the assumptions of an IV, then an interaction between risk factors has a causal interpretation. However, there is no way of distinguishing a purely statistical interaction from a mechanistic or biological interaction based on observational data. We therefore advise caution in the interpretation of interaction findings, as a statistical interaction can arise due to non-linearity in the effect of a risk factor, or because of the scale on which the outcome is measured (for example, an interaction may occur on the original scale, but not on a log-transformed scale). When considering an interaction between interventions, researchers can investigate whether there is an interaction between the interventions on the risk factor(s) as well as on the outcome. This may help reveal where any biological interaction may take place.

### Comparison with previous work

Previous work investigating interactions using IVs has been limited. A formal framework for defining interaction effects in the context of clinical trials was proposed by Blackwell [33], who used the language of principal stratification (compliance classes and monotonicity) to define local average interaction effects in a similar way to how local average causal effects (also called complier-averaged causal effects) are defined for single risk factors [34]. However, the principal stratification framework presupposes that risk factors are binary (or categorical) to assign compliance classes, whereas risk factors in Mendelian randomization are typically continuous. Additionally, the principal stratification framework presupposes a single binary instrumental variable, whereas Mendelian randomization investigations often use multiple genetic variants. There is therefore little practical advice in the literature on how to perform a factorial Mendelian randomization analysis.

### Limitations

There are several limitations to this work. We rely on the assumption that all genetic variants included in our analyses are valid IVs. If they are not, then estimates may be biased. Our recommendations rely on simulated data. Different choices for the parameters included in the simulation studies may have resulted in different conclusions. However, our findings were robust to different choices of parameters considered in this paper, they correspond to what we know about the theoretical properties of estimators, and similar conclusions were observed from the applied analysis. We have only considered interactions on an additive scale, although interactions could be considered on a multiplicative scale by log-transforming the outcome. Finally, we have not considered the impact of model misspecification on estimates. It would not be possible to perform simulation studies corresponding to all possible ways that model misspecification could occur, meaning that our recommendations cannot be proven to be optimal in all settings. We believe that we have chosen parameters and scenarios that are relevant to modern Mendelian randomization analyses.

### Conclusion

Overall, factorial Mendelian randomization is a promising technique for assessing interactions using genetic variants as instrumental variables. Our findings suggest that current applications of factorial Mendelian randomization based on a 2 × 2 analysis could be improved by better selection of genetic variants, and by better choice of analysis method.

Conflict of Interest: none declared

## Supplementary Material

In the Supplementary Material, we provide more detail on the two simulation studies and the applied example presented in the paper.

### Simulation study 1: interactions between risk factors

The two risk factors *X*_1_ and *X*_2_ were generated for *i* = 1, 2,…, 10 000 participants from the following data-generating model:

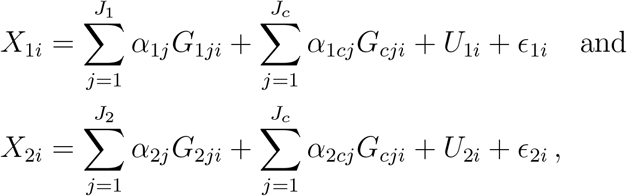

where ***G***_**1**_ and ***G***_**2**_ are the genetic variants associated with *X*_1_ and *X*_2_ respectively, and ***G***_***c***_ are the set of shared variants that are associated with both *X*_1_ and *X*_2_ (bold font represents vectors). The genotypes (0, 1 or 2) were generated independently from binomial distributions Bin(2, *MAF*_*j*_), where *MAF*_*j*_ represents the minor allele frequency (MAF) of the *j*^*th*^ genetic variant, and was drawn from a uniform distribution Unif(0.1, 0.5). ***α***_**1**_ and 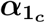 represent the effects of the genetic variants ***G***_**1**_ and ***G***_***c***_ on *X*_1_, and ***α***_**2**_ and 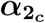 represent the effects of the genetic variants ***G***_**2**_ and ***G***_***c***_ on *X*_2_. The genetic associations were calculated so that ***G***_**1**_ and ***G***_***c***_, and ***G***_**2**_ and ***G***_***c***_, explained 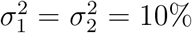 of the variance in *X*_1_ and *X*_2_ respectively. To ensure that each genetic variant explained the same amount of variation in the risk factor, we rearranged:

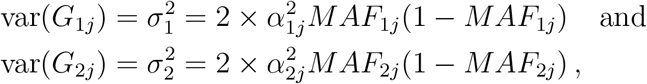

to calculate the genetic associations:

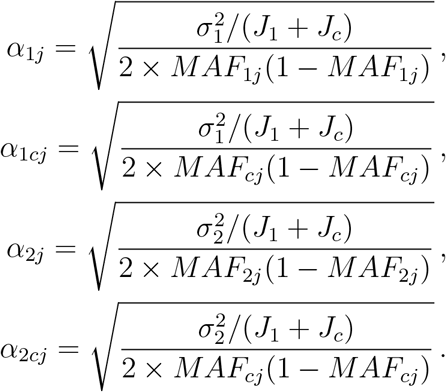

*U*_1_ and *U*_2_ represent the set of confounding variables of the *X*_1_ − *Y* and *X*_2_ − *Y* associations. To ensure the confounders explained 25% of the variation in the risk factors, *U*_1_ and *U*_2_ were drawn independently from a normal distribution 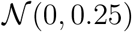. To fix the variances of *X*_1_ and *X*_2_ to one, the error terms *ϵ*_1_ and *ϵ*_2_ were generated independently from a normal distribution with mean zero, and variance:

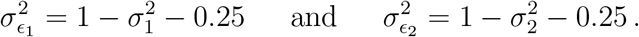

The outcome *Y* was generated from:

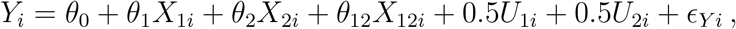

where *θ*_1_ and *θ*_2_ represent the main effects of *X*_1_ and *X*_2_ on *Y*, and *θ*_12_ represents the interaction effect of *X*_1_ and *X*_2_ on *Y*. *X*_12_ was generated by either: a) multiplying *X*_1_ and *X*_2_; or b) multiplying the mean centred values of the risk factors (*X*_1_ − *X̄*_1_) and (*X*_2_ − *X̄*_2_), where *X̄*_1_ and *X*̄_2_ are the mean values of *X*_1_ and *X*_2_. To ensure the risk factors and confounders explained less than a third of the variance in the outcome, the error term *ϵ*_*Y*_ was generated from a standard normal distribution 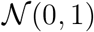.

Two-stage least squares regression models were fitted to either: a) the directly generated values of the risk factors (*X*_1_, *X*_2_, *X*_12_ = *X*_1_ × *X*_2_); or b) the mean centred values of the risk factors (*X*_1_ − *X̄*_1_, *X*_2_ − *X̄*_2_, *X*_12_ = (*X*_1_ − *X̄*_1_) × (*X*_2_ − *X̄*_2_)). When the risk factors were mean centred, the model estimated the marginal effects *θ*_1*M*_ and *θ*_2*M*_ of *X*_1_ and *X*_2_ on *Y*, otherwise *θ*_1_ and *θ*_2_ were estimated. For example, when there were no shared variants *J*_*c*_ = 0, the marginal effects were approximately:

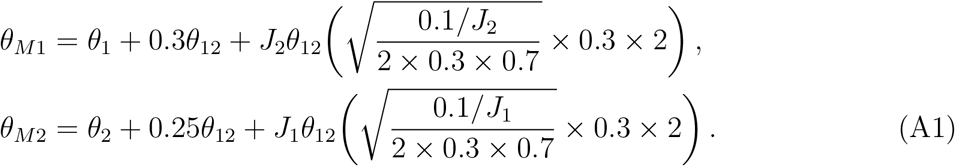

The genetic variants were either treated as individual IVs or as a single instrument in externally weighted gene scores 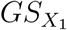 and 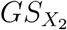 for *X*_1_ and *X*_2_. The external weights for the gene scores were based on an independent set of 10 000 individuals, and were produced from the same data generating model used for the main set of participants. The following four sets of genetic variants were used as IVs in separate two-stage least squares regression models:

- Method 1 – full set of interactions: the *J*_1_, *J*_2_ and *J*_*c*_ genetic variants used to generate *X*_1_ and *X*_2_, plus the unique interactions and quadratic terms of (***G***_**1**_ + ***G***_***c***_) × (***G***_**2**_ + ***G***_***c***_).
- Method 2 – reduced set of interactions: the *J*_1_, *J*_2_ and *J*_*c*_ genetic variants used to generate *X*_1_ and *X*_2_, plus the interactions from the product ***G***_**1**_ × ***G***_**2**_.
- Method 3 – continuous gene scores: the two weighted gene scores 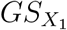 and 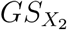, and their product 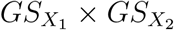.
- Method 4 – dichotomized gene scores: the two dichotomized gene scores, and their product.

Method 1 represents the oracle model as it includes all of the variables used in the data generating model, whereas Methods 2 to 4 are misspecified and their performance should be compared to Method 1. In Method 2, we have included a subset of the cross-terms between the genetic variants to create a more realistic scenario where the full set of relevant IVs are not included in the analysis. Method 3 considers the impact of including all of the genetic variants into two separate weighted gene scores, and finally, Method 4 considers the impact of dichotomizing the weighted gene scores.

Data were generated 10 000 times with *θ*_0_ = 0.2, *θ*_1_ = 0.3, *θ*_2_ = 0.2, and *θ*_12_ = 0.1, 0.3 and 0.5. Each risk factor was associated with (*J*_1_ + *J*_*c*_) = (*J*_2_ + *J*_*c*_) = 10 genetic variants, and the number of shared variants *J*_*c*_ was initially set to 0 to consider the scenario where none of the genetic variants were associated with risk factors (Table 1). The analyses were re-performed on the mean centred risk factors (Supplementary Table A1), and the number of shared variants was set to *J*_*c*_ = 1, 3, 5, 8 and 10 (Table 2). The following measurements were recorded for the estimates of *θ*_1_, *θ*_2_ and *θ*_12_: median estimate; standard deviation of estimates; median standard error of estimates; empirical power at the 5% significance level; and empirical coverage of the 95% confidence interval. The data were re-generated for 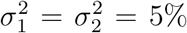 and 1%, for *J*_*c*_ = 0 (Supplementary Table A2) and *J*_*c*_ = 5 (Supplementary Table A3), and the analyses were re-performed on the directly generated values of the risk factors. Estimates of the F-statistic and conditional F-statistic for *X*_1_, *X*_2_ and *X*_12_ were recorded. The conditional F-statistic (also known as the Sanderson–Windmeijer F-statistic [1]) represents the strength of the IVs for the risk factors in a joint model, and is the relevant measure of instrument strength for a multivariable Mendelian randomization analysis [2].

**Supplementary Table A1:** Simulation study results for interactions between risk factors with no shared variants after centering the risk factors: median estimate, standard deviation (SD) of estimates, median standard error (SE), empirical power (%) to reject null at 5% significance, and empirical coverage (%) of 95% confidence interval. Note that centering changes the estimands for the main effect terms, not only the estimates – hence coverage is only displayed for the interaction term.

|  | Median | SD | Median SE | Power (%) | Coverage (%) |
| --- | --- | --- | --- | --- | --- |
| Methods 1 & 2 <sup>a</sup> – full set of interactions |  |  |  |  |  |
| $\theta_1 = 0.3$ | 0.4311 | 0.0327 | 0.0320 | 100.0 | - |
| $\theta_2 = 0.2$ | 0.3370 | 0.0328 | 0.0320 | 100.0 | - |
| $\theta_{12} = 0.1$ | 0.1101 | 0.0721 | 0.0718 | 33.7 | 94.6 |
| $\theta_1 = 0.3$ | 0.6679 | 0.0408 | 0.0320 | 100.0 | - |
| $\theta_2 = 0.2$ | 0.5823 | 0.0413 | 0.0320 | 100.0 | - |
| $\theta_{12} = 0.3$ | 0.3080 | 0.0722 | 0.0718 | 98.8 | 95.2 |
| $\theta_1 = 0.3$ | 0.9044 | 0.0527 | 0.0320 | 100.0 | - |
| $\theta_2 = 0.2$ | 0.8290 | 0.0528 | 0.0320 | 100.0 | - |
| $\theta_{12} = 0.5$ | 0.5073 | 0.0715 | 0.0718 | 100.0 | 95.2 |
| Method 3 – continuous gene scores |  |  |  |  |  |
| $\theta_1 = 0.3$ | 0.4178 | 0.0348 | 0.0343 | 100.0 | - |
| $\theta_2 = 0.2$ | 0.3234 | 0.0349 | 0.0343 | 100.0 | - |
| $\theta_{12} = 0.1$ | 0.1010 | 0.1113 | 0.1091 | 15.4 | 95.5 |
| $\theta_1 = 0.3$ | 0.6539 | 0.0424 | 0.0343 | 100.0 | - |
| $\theta_2 = 0.2$ | 0.5691 | 0.0431 | 0.0343 | 100.0 | - |
| $\theta_{12} = 0.3$ | 0.3000 | 0.1106 | 0.1091 | 77.5 | 95.8 |
| $\theta_1 = 0.3$ | 0.8906 | 0.0539 | 0.0343 | 100.0 | - |
| $\theta_2 = 0.2$ | 0.8165 | 0.0543 | 0.0343 | 100.0 | - |
| $\theta_{12} = 0.5$ | 0.4995 | 0.1107 | 0.1092 | 98.7 | 95.6 |
| Method 4 – dichotomized gene scores |  |  |  |  |  |
| $\theta_1 = 0.3$ | 0.4173 | 0.0438 | 0.0435 | 100.0 | - |
| $\theta_2 = 0.2$ | 0.3236 | 0.0438 | 0.0434 | 100.0 | - |
| $\theta_{12} = 0.1$ | 0.1022 | 0.1786 | 0.1720 | 8.0 | 95.9 |
| $\theta_1 = 0.3$ | 0.6538 | 0.0496 | 0.0435 | 100.0 | - |
| $\theta_2 = 0.2$ | 0.5687 | 0.0506 | 0.0435 | 100.0 | - |
| $\theta_{12} = 0.3$ | 0.2972 | 0.1777 | 0.1722 | 41.8 | 96.0 |
| $\theta_1 = 0.3$ | 0.8913 | 0.0597 | 0.0435 | 100.0 | - |
| $\theta_2 = 0.2$ | 0.8165 | 0.0603 | 0.0435 | 100.0 | - |
| $\theta_{12} = 0.5$ | 0.5002 | 0.1776 | 0.1718 | 80.7 | 96.1 |

<sup>a</sup>As there are no shared variants, methods 1 and 2 are equivalent.

**Supplementary Table A2:** Simulation study results for interactions between risk factors varying the amount of variance in the risk factors explained by the genetic variants, with no shared variants and an interaction effect *θ*_12_ = 0.3: mean F-statistic (F-stat), mean conditional F-statistic (CF-stat), median estimate, standard deviation (SD) of estimates, median standard error (SE), empirical power (%) to reject null at 5% significance, and empirical coverage (%) of 95% confidence interval.

|  | F-stat | CF-stat | Median | SD | Median SE | Power | Coverage |
| --- | --- | --- | --- | --- | --- | --- | --- |
| Variants explain 10% of the variance in risk factors: |  |  |  |  |  |  |  |
| Methods 1 & 2 <sup>a</sup> – full set of interactions |  |  |  |  |  |  |  |
| $\theta_1 = 0.3$ | 10.3 (0.6) | 2.1 (0.3) | 0.3043 | 0.0918 | 0.0910 | 91.0 | 95.0 |
| $\theta_2 = 0.2$ | 10.3 (0.6) | 2.1 (0.3) | 0.2034 | 0.0947 | 0.0945 | 57.9 | 95.5 |
| $\theta_{12} = 0.3$ | 8.1 (0.6) | 1.9 (0.2) | 0.3080 | 0.0722 | 0.0718 | 98.8 | 95.2 |
| Method 3 – continuous gene scores |  |  |  |  |  |  |  |
| $\theta_1 = 0.3$ | 364.2 (23.4) | 104.5 (25.6) | 0.2998 | 0.1359 | 0.1332 | 61.9 | 95.6 |
| $\theta_2 = 0.2$ | 364.5 (23.2) | 103.9 (25.3) | 0.2019 | 0.1405 | 0.1387 | 31.5 | 95.8 |
| $\theta_{12} = 0.3$ | 273.7 (22.4) | 97.8 (22.8) | 0.3000 | 0.1106 | 0.1091 | 77.5 | 95.8 |
| Method 4 - dichotomized gene scores |  |  |  |  |  |  |  |
| $\theta_1 = 0.3$ | 224.2 (17.7) | 41.9 (13.4) | 0.3039 | 0.2145 | 0.2074 | 32.1 | 95.8 |
| $\theta_2 = 0.2$ | 224.4 (17.7) | 41.7 (13.3) | 0.2047 | 0.2236 | 0.2164 | 15.2 | 96.2 |
| $\theta_{12} = 0.3$ | 168.2 (16.3) | 40.0 (12.4) | 0.2972 | 0.1777 | 0.1722 | 41.8 | 96.0 |
| Variants explain 5% of the variance in risk factors: |  |  |  |  |  |  |  |
| Methods 1 & 2 <sup>a</sup> – full set of interactions |  |  |  |  |  |  |  |
| $\theta_1 = 0.3$ | 5.4 (0.4) | 1.5 (0.2) | 0.3174 | 0.0931 | 0.0920 | 92.4 | 94.5 |
| $\theta_2 = 0.2$ | 5.4 (0.4) | 1.4 (0.2) | 0.2166 | 0.0957 | 0.0959 | 62.0 | 94.8 |
| $\theta_{12} = 0.3$ | 3.9 (0.4) | 1.2 (0.2) | 0.3087 | 0.0889 | 0.0888 | 92.8 | 95.0 |
| Method 3 – continuous gene scores |  |  |  |  |  |  |  |
| $\theta_1 = 0.3$ | 170.2 (15.5) | 25.4 (11.7) | 0.2988 | 0.2298 | 0.2121 | 29.9 | 96.9 |
| $\theta_2 = 0.2$ | 170.1 (15.7) | 25.2 (11.5) | 0.1985 | 0.2421 | 0.2237 | 13.8 | 96.9 |
| $\theta_{12} = 0.3$ | 109.4 (13.3) | 23.8 (10.4) | 0.3020 | 0.2458 | 0.2276 | 26.7 | 96.9 |
| Method 4 - dichotomized gene scores |  |  |  |  |  |  |  |
| $\theta_1 = 0.3$ | 107.3 (12.2) | 10.7 (6.7) | 0.2970 | 3.928 | 0.3367 | 12.6 | 98.9 |
| $\theta_2 = 0.2$ | 106.9 (12.0) | 10.6 (6.6) | 0.1948 | 3.804 | 0.3551 | 5.4 | 98.7 |
| $\theta_{12} = 0.3$ | 68.8 (10.2) | 10.2 (6.1) | 0.3033 | 4.065 | 0.3654 | 10.8 | 98.8 |
| Variants explain 1% of the variance in risk factors: |  |  |  |  |  |  |  |
| Methods 1 & 2 <sup>a</sup> – full set of interactions |  |  |  |  |  |  |  |
| $\theta_1 = 0.3$ | 1.8 (0.2) | 1.4 (0.2) | 0.3681 | 0.0910 | 0.0901 | 97.7 | 88.4 |
| $\theta_2 = 0.2$ | 1.8 (0.2) | 1.4 (0.2) | 0.2670 | 0.0930 | 0.0930 | 81.4 | 88.6 |
| $\theta_{12} = 0.3$ | 1.4 (0.2) | 1.0 (0.1) | 0.3029 | 0.0971 | 0.0972 | 86.4 | 95.4 |
| Method 3 – continuous gene scores |  |  |  |  |  |  |  |
| $\theta_1 = 0.3$ | 29.5 (6.4) | 1.9 (2.9) | 0.2854 | 29.26 | 0.8411 | 2.8 | 99.9 |
| $\theta_2 = 0.2$ | 29.4 (6.4) | 1.9 (2.8) | 0.1883 | 31.58 | 0.9203 | 1.0 | 99.9 |
| $\theta_{12} = 0.3$ | 12.3 (4.1) | 1.6 (2.1) | 0.3185 | 52.32 | 1.537 | 0.7 | 100.0 |
| Method 4 - dichotomized gene scores |  |  |  |  |  |  |  |
| $\theta_1 = 0.3$ | 19.1 (5.1) | 1.6 (2.8) | 0.2992 | 123.8 | 1.063 | 1.9 | 99.9 |
| $\theta_2 = 0.2$ | 19.0 (5.0) | 1.5 (2.4) | 0.1930 | 217.5 | 1.163 | 0.6 | 100.0 |
| $\theta_{12} = 0.3$ | 8.1 (3.3) | 1.2 (1.7) | 0.3121 | 347.4 | 1.933 | 0.3 | 100.0 |

<sup>a</sup>As there are no shared variants, methods 1 and 2 are equivalent.

**Supplementary Table A3:**
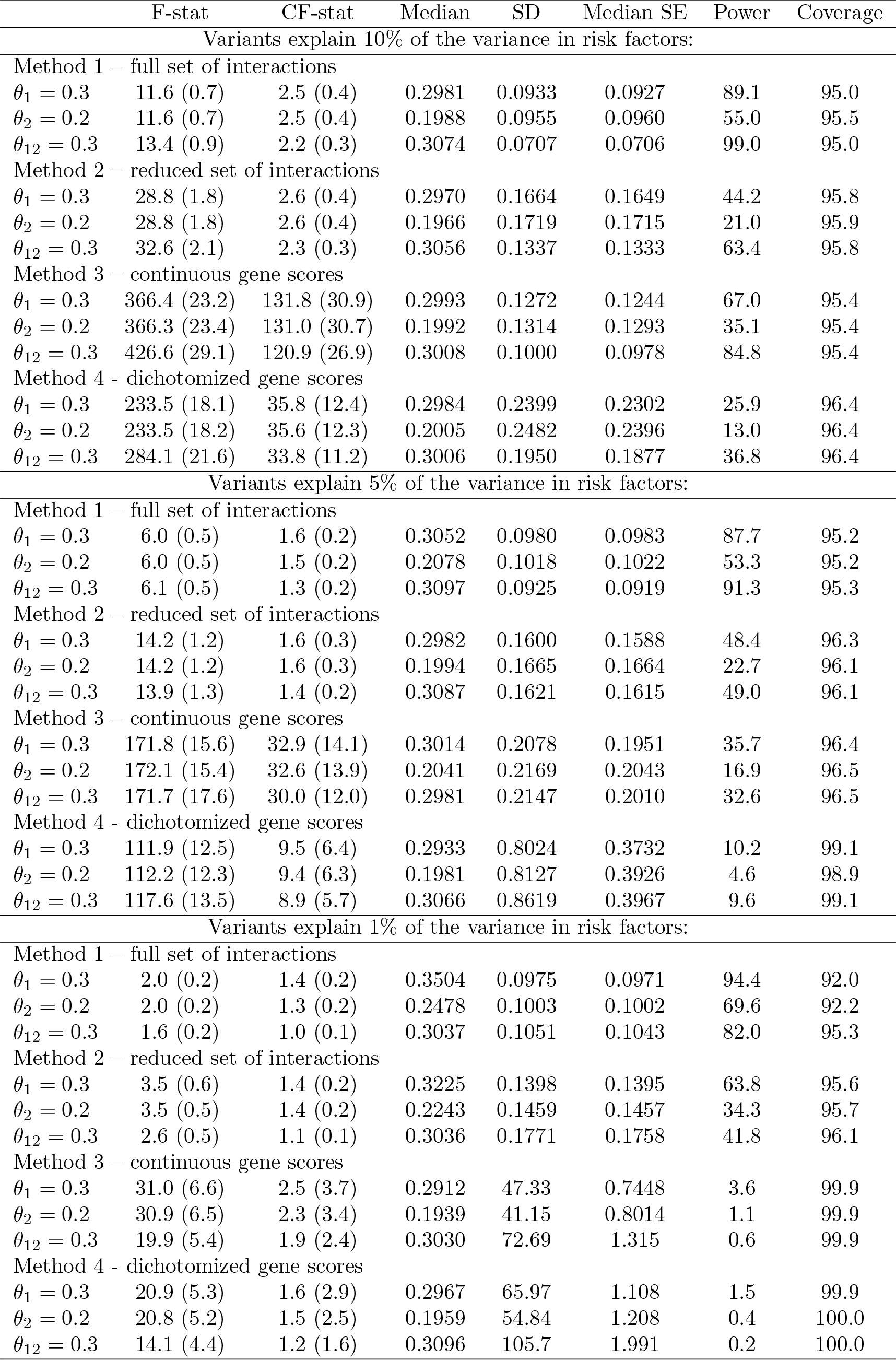
Simulation study results for interactions between risk factors varying the amount of variance in the risk factors explained by the genetic variants, with 5 shared variants and an interaction effect *θ*_12_ = 0.3: mean F-statistic (F-stat), mean conditional F-statistic (CF-stat), median estimate, standard deviation (SD) of estimates, median standard error (SE), empirical power (%) to reject null at 5% significance, and coverage (%) of 95% confidence interval.

### Simulation study 2: interactions between interventions

Using the same notation defined in the first simulation study, the risk factor *X* was generated for *i* = 1, 2*,…,* 10 000 participants from the following data generating model:

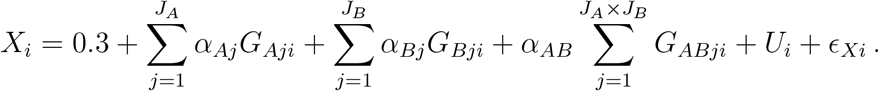

We assume that the two gene regions are distinct, and the genetic variants ***G***_***A***_ and ***G***_***B***_ are not in linkage disequilibrium. The genotypes were generated independently from binomial distributions Bin(2, *MAF*_*j*_), where *MAF*_*j*_ represents the MAF for the *j*^*th*^ genetic variant. *MAF*_*j*_ was drawn from a uniform distribution 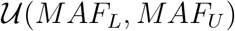, where the value of *MAF*_*L*_ and *MAF*_*U*_ were either taken as 0.4 and 0.5 (common variants), or 0.1 and 0.2 (uncommon variants). We assumed that the interaction effect *α*_*AB*_ was constant across the *J*_*A*_ × *J*_*B*_ product terms for simplicity.

The approximate proportion of variance explained in *X* by ***G***_***A***_ 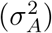 and ***G***_***B***_ 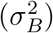 varied between scenarios. As before, the genetic associations ***α***_***A***_ and ***α***_***B***_ were calculated by rearranging the formula for the variance of the genetic variants to ensure the amount of variance explained by each variant was the same:

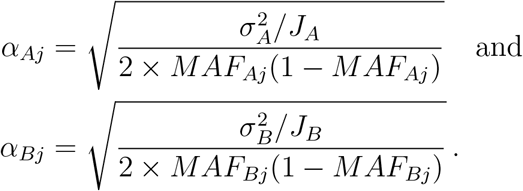

The confounders *U* were drawn from 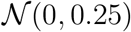, and the error term *ϵ*_*X*_ was generated from 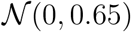. The outcome *Y* was generated from:

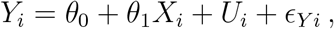

where *θ*_1_ represents the causal effect of *X* on *Y*, and the error term *ϵ*_*Y*_ was generated from a standard normal distribution 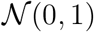. The data was generated 10 000 times under the following scenarios:

- Scenario 1: 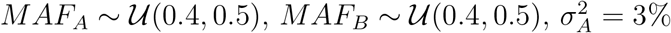 and 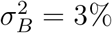
- Scenario 2: 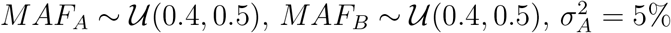 and 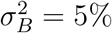
- Scenario 3: 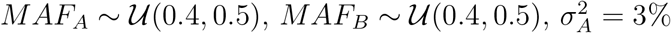 and 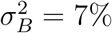
- Scenario 4: 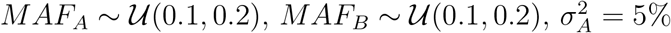 and 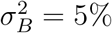
- Scenario 5: 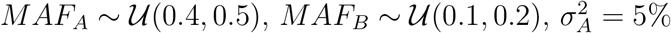 and 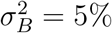
- Scenario 6: 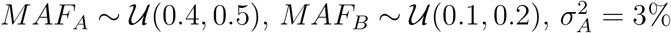 and 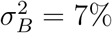

with *J*_*A*_ = *J*_*B*_ = 3, *θ*_0_ = 0.2, *θ*_1_ = 0.1, and *α*_*AB*_ = 0.1, 0.3 and 0.5. The above scenarios were selected to consider the impact of varying the MAF and the amount of variance in the risk factor explained by the genetic variants had on the performance of the method.

For each scenario, optimal weighted gene scores *GS*_*A*_ and *GS*_*B*_ were generated for each gene region, where the external weights were produced from an independent set of 10 000 individuals from the same data-generating model used for the main set of participants. The two gene scores were dichotomized at their median values to create two binary variables. The outcome was then regressed against: a) the two continuous gene scores and their product; and the dichotomized gene scores and their product. The following measurements were recorded for the estimate of the interaction effect between the gene scores on the outcome: median estimate; standard deviation of estimates; median standard error; and empirical power at the 5% significance level.

### Applied example: the effects of BMI and alcohol on systolic blood pressure

UK Biobank is a prospective, population-based cohort consisting of approximately 500,000 participants aged between 40 and 69 years at baseline living in the UK. Extensive baseline characteristics were collected at recruitment, including lifestyle factors, sociodemographic information, and physical attributes. For the analysis, we considered 367,643 unrelated participants of European descent who passed data quality control measures and had genetic data [3].

Body mass index (BMI, kg/m^2^) and systolic blood pressure (SBP, mmHg) were measured at baseline when participants attended the assessment centre. Information on baseline alcohol consumption was obtained from a touchscreen questionnaire which included questions on alcohol drinking status, frequency of alcohol consumption, and beverage type. The responses to the amount of alcohol drank and beverage type were used to create a continuous variable that represented alcohol consumption in units per day. To adjust for blood pressure medication, 15 mmHg was added to SBP for individuals who reported to be on blood pressure lowering medication [4]. Individuals were dropped from the analysis if they had missing data on BMI, SBP, alcohol consumption, or relevant genetic variants. The final sample size was 291,781.

We used the 77 genome-wide significant variants from a meta-analysis by the Genetic Investigation of ANthropometric Traits (GIANT) consortium in participants of European ancestry to act as IVs for BMI [5]. For alcohol, we identified 10 genetic variants in the *ADH1B* gene region that have been shown to be associated with alcohol consumption [6]. The genetic variants used as IVs for BMI and alcohol consumption were cross-referenced to check for any overlap. BMI was regressed separately against each of the 10 alcohol variants, and alcohol consumption was regressed against each of the 77 BMI variants. All models were adjusted for gender, age, and the first ten genomic principal components.

Internally-weighted gene scores were created for BMI based on the 77 genetic variants (*GS*_*BMI*_), and for alcohol consumption based on the 10 genetic variants (*GS*_*AC*_), and these gene scores were dichotomized at their median values to create two binary variables. A separate binary variable was generated using the rs1229984 variant only, where participants were either considered to have: a) a low alcohol consumption if they were homozygous or heterozygous for the alcohol-decreasing allele; or b) a high alcohol consumption if they were homozygous for the alcohol-increasing allele (as in the paper by Carter *et al.* [7]). Using these binary variables, the following groups of participants were created:

- Low BMI, low alcohol consumption: *GS*_*BMI*_ ≤ *med*(*GS*_*BMI*_) and *GS*_*AC*_ ≤ *med*(*GS*_*AC*_) or was homozygous or heterozygous for the alcohol decreasing allele for the rs1229984 variant,
- High BMI, low alcohol consumption: *GS*_*BMI*_ > *med*(*GS*_*BMI*_) and *GS*_*AC*_ ≤ *med*(*GS*_*AC*_) or was homozygous or heterozygous for the alcohol decreasing allele for the rs1229984 variant,
- Low BMI, high alcohol consumption: *GS*_*BMI*_ ≤ *med*(*GS*_*BMI*_) and *GS*_AC_ > *med*(*GS*_*AC*_) or was homozygous for the alcohol increasing allele for the rs1229984 variant, and
- High BMI, high alcohol consumption: *GS*_*BMI*_ ≤ *med*(*GS*_*BMI*_) and *GS*_*AC*_ > *med*(*GS*_*AC*_) or was homozygous for the alcohol increasing allele for the rs1229984 variant.

The above criteria created four groups of participants based on the dichotomized gene scores for BMI and alcohol consumption, and another four groups based on the dichotomized gene score for BMI and the rs1229984 variant. The numbers of participants, and the mean and standard deviation of BMI, alcohol consumption, and SBP were recorded for each group.

Two-stage least squares regression models of SBP were fitted to BMI, alcohol consumption, and the product of BMI and alcohol consumption. The following sets of IVs were considered:

- Method 1: the 77 variants for BMI and 10 variants for alcohol consumption, plus 770 cross-terms between the two sets of variants.
- Method 2: the continuous gene scores *GS*_*BMI*_ and *GS*_*AC*_, plus their product *GS*_*BMI*_ × *GS*_*AC*_.
- Method 3: the dichotomized gene scores of *GS*_*BMI*_ and *GS*_*AC*_, plus their product.

The models were refitted excluding all of the variants for alcohol consumption apart from the lead rs1229984 variant. All models were adjusted for gender, age, and the first ten genomic principal components. For each model, the estimate and standard error of the interaction term was recorded with its p-value. In total, six two-stage least squares regression models were fitted to the dataset, and all of the models were adjusted for age, gender and the first 10 genomic principal components. The F-statistic and the Sanderson–Windmeijer conditional F-statistic were estimated for each set of IVs with respect to BMI, alcohol consumption, and the product of BMI and alcohol consumption (Supplementary Table A4).

**Supplementary Table A4:** F-statistics (F-stat) and conditional F-statistics (CF-stat) for applied example.

|  | Method 1 |  | Method 2 |  | Method 3 |  |
| --- | --- | --- | --- | --- | --- | --- |
|  | F-stat | CF-stat | F-stat | CF-stat | F-stat | CF-stat |
| 10 variants for alcohol: |  |  |  |  |  |  |
| BMI | 6.8 | 1.3 | 1662.8 | 21.1 | 1054.1 | 7.0 |
| Alcohol consumption | 2.4 | 1.1 | 268.0 | 20.9 | 55.6 | 6.9 |
| Product term | 2.4 | 1.1 | 298.6 | 21.0 | 73.2 | 6.9 |
| rs1229984 for alcohol: |  |  |  |  |  |  |
| BMI | 32.8 | 1.3 | 1654.9 | 17.2 | 1066.8 | 13.5 |
| Alcohol consumption | 7.7 | 1.2 | 245.1 | 17.1 | 241.6 | 13.4 |
| Product term | 7.9 | 1.2 | 267.7 | 17.1 | 266.5 | 13.4 |

